# Goblet cell LRRC26 regulates BK channel activation and protects against colitis in mice

**DOI:** 10.1101/2020.10.23.341396

**Authors:** Vivian Gonzalez-Perez, Pedro L. Martinez-Espinosa, Monica Sala-Rabanal, Nikhil Bharadwaj, Xiao-Ming Xia, Albert C. Chen, David Alvarado, Jenny K. Gustafsson, Hongzhen Hu, Matthew A. Ciorba, Christopher J. Lingle

## Abstract

Goblet cells (GCs) are specialized cells of the intestinal epithelium contributing critically to mucosal homeostasis. One of the functions of GCs is to produce and secrete MUC2, the mucin that forms the scaffold of the intestinal mucus layer coating the epithelium and separates the luminal pathogens and commensal microbiota from the host tissues. Although a variety of ion channels and transporters are thought to impact on MUC2 secretion, the specific cellular mechanisms that regulate GC function remain incompletely understood. Previously, we demonstrated that leucine-rich-repeat-containing protein 26 (LRRC26), a known regulatory subunit of the Ca^2+^-and voltage-activated K^+^ channel (BK channel), localizes specifically to secretory cells within the intestinal tract. Here, utilizing a mouse model in which MUC2 is fluorescently tagged allowing visualization of single GCs in intact colonic crypts, we show that murine colonic GCs have functional LRRC26-associated BK channels. In the absence of LRRC26, BK channels are present in GCs, but are not activated at physiological conditions. In contrast, all tested MUC2-negative cells completely lacked BK channels. Moreover, LRRC26-associated BK channels underlie the BK channel contribution to the resting transepithelial current across mouse distal colonic mucosa. Genetic ablation of either LRRC26 or BK-pore forming α-subunit in mice results in a dramatically enhanced susceptibility to colitis induced by dextran sodium sulfate (DSS). These results demonstrate that normal potassium flux through LRRC26-associated BK channels in GCs has protective effects against colitis in mice.

**Significance:** A primary function of goblet cells (GCs) of the intestinal epithelium is to generate a protective mucus layer lining the intestinal lumen. GC dysfunction is linked to Inflammatory Bowel Disease (IBD). GC mucus secretion is thought to be dependent on contributions of an ensemble of anion and cation fluxes, although understanding remains limited. Here, it is shown in mouse colon that the Ca^2+^- and voltage-dependent BK-type K^+^ channel, specifically in association with the LRRC26 regulatory subunit, plays a critical role in normal GC function, protecting mice against chemically-induced colitis. The results demonstrate that normal K^+^ fluxes mediated by LRRC26-containing BK channels are required for normal GC function, potentially providing insights into the potential role of BK channels in IBD.

## Introduction

The colonic epithelium is composed of a single layer of heterogeneous cells, covered by mucus, that separate the luminal contents from host tissues. Acting both in concert and individually, the diverse cells comprising the epithelial layer play the functions of protection (1), sensation (2, 3), transport of substances (4, 5), and repair (6). Colonic epithelial cells belong to three lineages: absorptive enterocytes, enteroendocrine cells, and goblet cells (GCs). The colonic epithelium is morphologically organized into repeating units called crypts of Lieberkühn where stem cells located at the base of the crypts divide and successively differentiate into the mature lineages as they migrate toward the crypt surface (7). Many of the key specialized functions of epithelial cells are, in part, defined by proteins involved in ion transport, located either on their luminal or basolateral membrane. Thus, among different gastrointestinal epithelial cells, ion channels, carriers, exchangers, and pumps work in concert to define a variety of essential functions: 1) solute and electrolyte absorption and secretion in absorptive enterocytes (reviewed in (5, 8); 2) environment sensation and serotonin secretion by enteroendocrine cells (2, 9); and 3) mucus secretion by GCs and subsequent mucus maturation into the protective layer covering the epithelial surface (10–12). In contrast to the ionic transport entities involved in electrolyte and fluid transport or environment sensation, ionic transport in GCs and its implications in GC physiology are much less understood. Here, we address the role of the Ca^2+^- and voltage-activated K^+^ channel (BK channel) in GCs.

GCs play two primary roles: one related to the maintenance of the mucosal barrier (reviewed in (1, 13)) and one related with the mucosal immune homeostasis (reviewed in (14, 15)). The role of GCs in barrier maintenance consists in generation of the mucus layer lining the intestinal lumen. One way GCs carry out this role is by secreting MUC2, the gel-forming mucin that forms the scaffold of the mucus layer separating luminal pathogens and commensal microbiota from the epithelial surface (11, 12, 15, 16). This separation is critical as has been demonstrated in both animal models and humans: mouse models with deficient mucus layer generation develop spontaneous colitis (16, 17), whereas a more penetrable mucus layer has been observed in patients with Ulcerative Colitis (18, 19). The constant replenishment of the mucus layer involves MUC2 exocytosis from GCs, and subsequent maturation (hydration and expansion) of the secreted MUC2 to form the gel-like mucus coating the epithelium (15). Both exocytosis and maturation of MUC2 are highly dependent on anion and K^+^ transport (10–12, 20). It has been proposed that mucin exocytosis in colon requires activities of the Na^+^/K^+^/2Cl^-^ cotransporter (NKCC1) (20, 21), and also anion and K^+^ channels whose identities are still unclear (20). It is also not clearly known whether specific ionic conductances are intrinsic to GCs or are located in the surrounding absorptive enterocytes. Although several types of K^+^ channels including K_Ca_3.1, Kv7.1 and BK channels have been found in colonic epithelial cells (22–27), to what extent any are specifically associated with GCs or critical to their function remains unclear. To date, most functional studies about colonic K^+^ channels have focused on their roles in electrolyte and fluid secretion/absorption of the whole colon, whereas the cellular events relating K^+^ channels to specific roles in GC function are still poorly understood.

Among colonic epithelial K^+^ channels, the BK channel (also known as K_Ca_1.1), the Ca^2+^-and voltage-activated K^+^ channel of high conductance, has been proposed to be the main component of colonic K^+^ secretion to the lumen (28–30). BK channels are homotetramers of the pore-forming BKα subunit, but can also contain tissue-specific regulatory subunits that critically define the functional properties of the channel (31). BK channels composed exclusively of the pore-forming BKα subunit are unlikely to be activated at the physiological conditions of epithelial cells and, as a consequence, the molecular properties of colonic BK channels that would allow them to contribute to colonic ion transport remain unclear. Recently, we established that the leucine-rich-repeat-containing protein 26 (LRRC26), a BK regulatory gamma subunit, is specifically expressed in secretory epithelial cells, including GCs of the gastrointestinal tract (32). When LRRC26 is present in a BK channel complex, the resulting channel activates near normal resting physiological conditions, even in the absence of any elevation of intracellular Ca^2+^(33).

In the present study, we have specifically probed the role of BK channels in cells of the colonic epithelium and examined the impact of deletions of either the BK α subunit or LRRC26 on colonic function. Here, through recordings from identified GCs in intact colonic crypts, we show that LRRC26-associated BK channels contribute the major K^+^ current in mouse colonic GCs. Furthermore, the LRRC26-containing BK channels are activated near −40 mV even in the absence of intracellular Ca^2+^. In contrast, in identified GCs from *Lrrc26^-/-^* mice, BK current is present, but it is only activated at membrane potentials unlikely to ever occur physiologically. Surprisingly, all colonic epithelial MUC2-negative cells sampled completely lack functional BK channels. To establish that the LRRC26-containing BK channels contribute to normal K^+^ fluxes in intact colon tissue, we show that the transepithelial current across distal colon at rest has a component dependent on LRRC26-associated BK channels, which is absent when either BKα or LRRC26 is genetically deleted. Moreover, the genetic ablation of either LRRC26 or BK channel results in a dramatically enhanced susceptibility to colitis induced by dextran sodium sulfate (DSS). Overall, our results suggest that normal potassium flux through LRRC26-associated BK channels in GCs has protective role against development of colitis.

## Results

### LRRC26 is specifically expressed in epithelial goblet cells in mouse colon

*Lrrc26^-/-^* mice, as previously described, have the entire genomic sequence of *Lrrc26* replaced with a cassette including *LacZ* gene (32). Using the β-GAL enzymatic activity as reporter, *Lrrc26* promoter activity is primarily observed in secretory epithelial cells including those in the gastrointestinal tract (32). Here, we examined the *Lrrc26* promoter activity in mouse colon in more detail. After incubation with X-Gal substrate, longitudinal sections of *Lrrc26^-/-^* and *WT* colons dissected from littermate mice identify only a subset of epithelial cells as positive for *Lrrc26* promoter activity (Fig. 1A-B, Fig. S1). Positive cells are found at all levels of the crypts and also in the luminal brush border surface. Positive cells in the upper most luminal third of the crypts exhibit a morphology typical of differentiated GCs (34). The number of *Lrrc26* positive cells gradually increases moving from the proximal towards the distal end of the colon, with the larger number of blue-stained cells per crypt in the 2/3 most distal segments (Fig. S1) and the highest number per crypt in the region closer to the anorectal junction. Blue staining was not observed in *lamina propria* or in the smooth muscle cell layer even after varying fixation conditions or using longer substrate exposure (up to 24 h) (Fig. S2).

**Figure 1.**
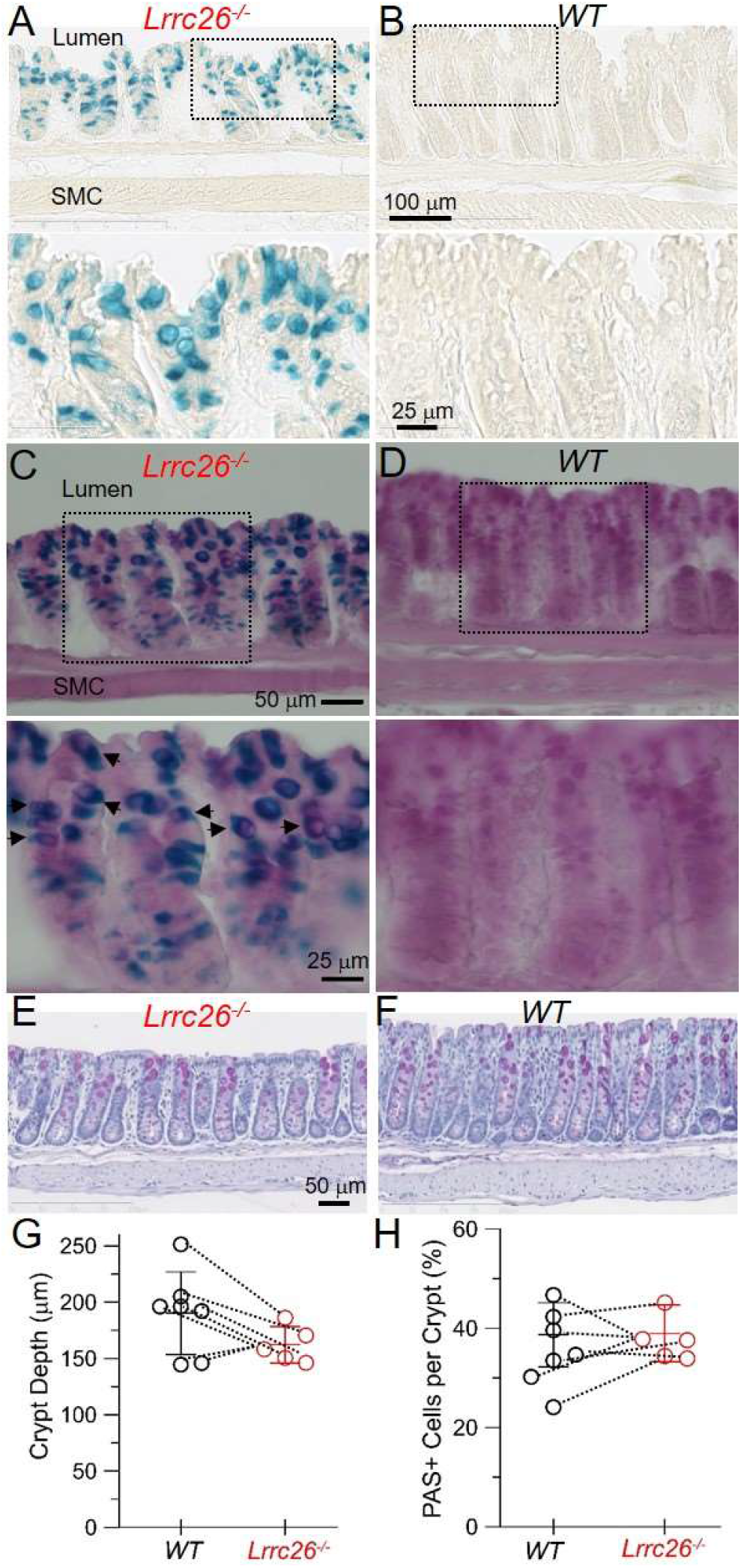
In mouse colon, LRRC26 is specifically expressed in epithelial goblet cells. Representative images of glutaraldehyde-fixed frozen 20 micron sections of *Lrrc26^-/-^* and *WT* distal colons from littermates, stained side-by-side with X-Gal substrate for 1 h (**A-B**) or X-Gal for 1 h, followed by PAS staining (**C-D**). Regions outlined in the boxes (dotted lines) are shown at higher magnification below with arrowheads identifying cells positive for both *Lrrc26*-promoter activity and PAS staining. Images were taken at 0.5 cm from the anorectal junction. SMC, smooth muscle cell layer. **E-H**, Morphological comparison between *Lrrc26^-/-^* and *WT* distal colon. Representative images of paraformaldehyde-fixed paraffin embedded 5 micron sections of distal third of colons from *Lrrc26^-/-^* (**E**) and *WT* (**F**) littermates (taken at 1 cm from the anorectal junction), stained with PAS and counterstained with hematoxylin to identify nuclei. Comparison of crypt depth (**G**) and relative abundance of GCs per crypt (**H**) between genotypes (mean ± SD). Data plotted correspond to 5 sets of littermates, both sexes included, dissected at approximately same age (8-12 weeks). Dotted lines connected results from siblings. Sets of values were compared with two-tailed Student’s t-test, P=0.0724 for crypt depth, P=0.3154 for GC abundance.

In order to verify that *Lrrc26* positive cells are GCs, we used Periodic Acid-Schiff (PAS) staining. PAS stains polysaccharide-rich cells and, therefore, readily identifies GC mucin granules (35). PAS is the classical method to histologically identify intestinal secretory cells which in colon are mainly GCs. X-Gal/PAS staining of frozen sections mildly fixed with either glutaraldehyde (Fig. 1C-D) or paraformaldehyde (Fig. S2) reveals that essentially all β-Gal positive cells in the brush border or the ¾ outermost part of the crypt were also PAS positive. A few blue-stained cells located at the base of the crypts in the most distal third of colon exhibited no appreciable PAS staining (Fig. S2A, C), perhaps suggesting that *Lrrc26* may also be expressed in stem cells. Overall, our results indicate that, in mouse colon, *Lrrc26* is expressed in epithelial GCs at all stages of differentiation. These results are consistent with a single cell screening of mouse intestinal epithelial cells where *Lrrc26* has been identified as a GC marker (36).

We next examined whether KO of LRRC26 has consequences on colonic morphology and epithelial cell composition by comparing *Lrrc26^-/-^* and *WT* colon histologic sections after thorough tissue fixation and staining with hematoxylin/eosin (Fig. S3) or PAS (Fig. 1E-F, Fig. S3). We analyzed the crypt metrics and GC abundance in the most distal third. Our results indicate that there are no differences between *Lrrc26^-/-^* and *WT* littermates in terms of crypt depth or abundance of GCs (Fig. 1G-H).

### Goblet cell BK channels contain LRRC26, endowing them with ability to be activated at resting physiological conditions

We have previously found that BK channels in acinar cells of lacrimal and salivary glands contain LRRC26 subunits (32). In colonic mucosa, the presence of BK channels has been documented for several species including humans (22, 27, 30, 37, 38), although the cell type(s) where it is expressed remain unclear. Using patch clamp, we directly tested whether colonic GCs have functional BK channels with properties consistent with the presence of LRRC26. In order to unambiguously identify the GCs, we used a transgenic mouse in which a mCherry-tag is attached to MUC2, the main mucin produced and secreted by gastrointestinal GCs (39). Thus, fluorescent cells are MUC2 positive cells, indicative of GCs. By mating this mouse with our *Lrrc26^-/-^* mouse model, we also generated *Lrrc26^-/-^* mice with fluorescent GCs.

We recorded the whole cell current in individual fluorescent cells of intact crypts freshly isolated from the most distal third colonic segment of *WT* (Fig. 2A-B) or *Lrrc26^-/-^* mice (Fig. 2C-D). We used solutions containing physiological Na^+^ and K^+^ gradients, but with glutamate replacing most of Cl’ in the solutions to minimize the contribution of anion currents, and an intracellular solution containing a Ca^2+^ concentration buffered to 250 nM, similar to conditions that we have previously utilized for patch clamping salivary gland cells (32). Our results show that, in fluorescent cells (*i.e*., GCs), depolarizing voltage steps evoke a time-dependent K^+^ current typical of BK channels, but with markedly shifted range of current activation between *WT* and *Lrrc26^-/-^* cells (Fig. 2B, D, H). In *WT* GCs, the K^+^ current begins to be activated at a membrane potential around - 40 mV with an activation curve for individual cells having on average a voltage of half activation of conductance (V_h_) = 13.4 ± 3.2 mV. In contrast, in *Lrrc26^-/-^* GCs, depolarizing steps up to +100 mV (red traces in Fig. 2B, D) barely activate any K^+^ current. Further depolarization does evoke a large K^+^ current, with the features of BK, with an activation curve having on average a V_h_ = 158.7 ± 5.1 mV for individual *Lrrc26^-/-^* GCs. Both *WT* and *Lrrc26^-/-^* GC-K^+^ currents were pharmacologically confirmed as BK (Fig. 2E-G). *WT* currents were reversely blocked by bathing the cells with 5 mM of tetraethyl ammonium (TEA), and both *WT* and *Lrrc26^-/-^* currents were completely inhibited by 200 nM of paxilline, a BK-specific blocker (40, 41). A Boltzmann fit to the averaged fractional conductance-voltage (G-V) curve generated from tail currents (Fig. 2H) yielded a V_h_ =13.5 mV and *z* = 1.13 *e_0_* for *WT*-GCs, while a V_h_ = 158.5 mV and z=1.05 *e_0_* for *Lrrc26^-/-^* GCs. These results indicate that genetic ablation of LRRC26 results in a 145 mV rightward shift of the activation curve of GC-BK channels (Fig 2H). The magnitude of the effect of LRRC26 on the activation of BK channels is consistent with the known effects of LRRC26 on heterologously expressed BK channels (42) or on native BK channels from salivary glands (32). The impact of LRRC26 is so pronounced that GC-BK channels lacking LRRC26 are hardly open and likely non-functional under physiological conditions. Overall, these results indicate that mouse colonic GCs have LRRC26-associated BK channels whose gating range is suitable to permit K^+^ efflux near normal physiological conditions, but only when LRR26 is present. Furthermore, the results show that BK channels are the primary K^+^ current present in GCs under these conditions.

**Figure 2.**
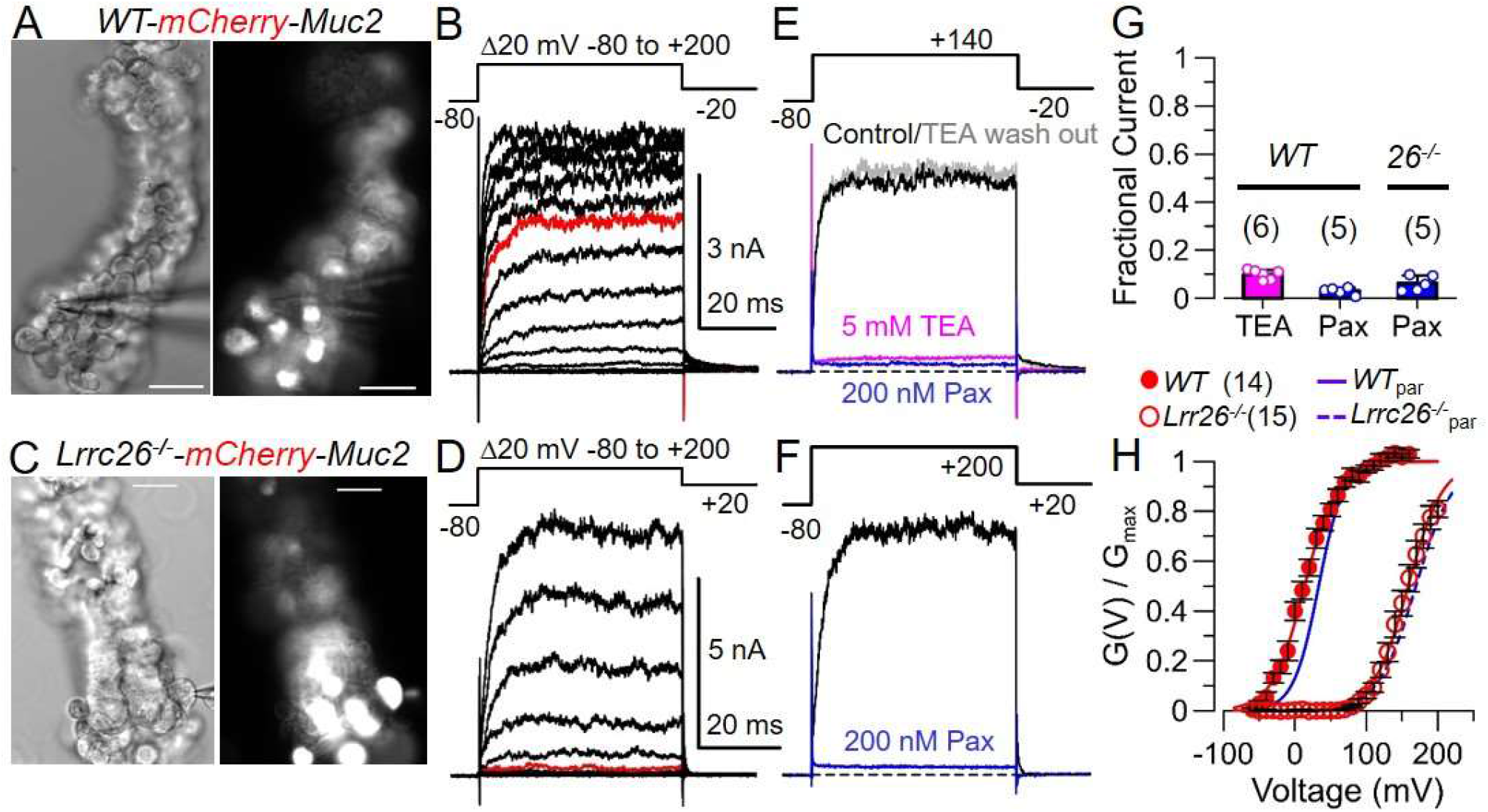
BK currents in GCs exhibit properties consistent with coassembly with LRRC26 permitting channel activation at resting physiological conditions. **A**, Bright field and fluorescent images of a colonic crypt isolated from *WT* mouse (crypt oriented with opening facing downward), with the recording pipette attached to the fluorescence-identified GC whose current is shown in (**B**) and (**E**); scale bars are 10 μm. **B,** Representative whole-cell K^+^ currents evoked by the depicted voltage protocol (top) in *WT*-GCs. **C-D**, Similar experiment as in A-B performed in intact colonic crypt isolated from *Lrrc26^-/-^* mouse showing that knock out of LRRC26 removes K^+^ currents activated by depolarizations up to +100 mV (red traces). Dotted lines shows zero current level. **E**, K^+^ current inhibition at +140 mV (maximal BK activation), in the same *WT* GC shown in (A-B), produced by extracellular 5 mM TEA (magenta), after TEA washout (gray), and finally with 200 nM paxilline (blue). **F**, K^+^ current inhibition in the same *Lrrc26^-/-^* GC shown in (C-D) by 200 nM paxilline (blue). Note the much larger depolarizing step (+200 mV) required to fully activate BK current in the *Lrrc26^-/-^* GC. **G,** Summary of pharmacological experiments confirming original current as mostly BK current (bars plot mean ±SD with points plotting measurements from individual cells). **H,** Voltagedependence of activation of BK conductance obtained from recordings as in (B) and (D). Fractional Conductance-Voltage (G(V)/G_max_) curves were generated from the tail conductances (G(V)), normalized to the maximum conductance (G_max_) from a Boltzmann fit of each individual cell, and averaged (± SEM). From Boltzmann fit to individual cells, for *WT* GCs (n=14), mean V_h_ = 13.4 ± 3.2 mV and *z* = 1.23 ± 0.05 e_0_; for *Lrrc26^-/-^* GCs (n=15), mean V_h_ = 158.7 ± 5.1 mV and *z* = 1.25 ± 0.06 *e_0_*. Fit of averaged G(V)/G_max_ curves yielded similar values: *WT* GCs, V_h_ = 13.5 mV, *z* = 1.13 *e_0_*; *Lrrc26^-/-^* GCs, V_h_ = 158.5 mV, *z* = 1.1. As reference, activation curves of LRRC26-associated BK or BK (lacking LRRC26) from parotid cells are shown in blue.

### Colonic epithelial MUC2-negative cells tested lack BK channels

Next, we examined whether colonic crypt cells other than GCs have functional BK channels. Using the same preparation and ionic conditions as above, we patch clamped non-fluorescent cells from *WT* mCherry-Muc2 transgenic mice. In non-fluorescent cells (i.e., MUC2-negative cells), which in the mouse distal colon are predominantly columnar enterocytes, the same voltage protocol applied previously only evokes a small, time-independent current (Fig. 3A-B) that linearly increases with the magnitude of the depolarization (Fig. 3C). This voltage-independent current has, on average, a reversal potential (E_rev_) of −35.2 ± 14.4 mV (mean ± SD) for individual cells (Fig. 3D). The fact that, in most of MUC2-negative cells sampled, this current reverses at a potential apart from the theoretical K^+^ reversal potential (E_K+_) suggests that it might result from the contribution of several conductances that may include both K^+^-selective and Na^+^-selective entities. The large variability of individual E_rev_ observed (Fig. 3D) might reflect the existence of different subsets of MUC2-negative cells having different sets of conductances. K^+^ channels previously identified in colonic epithelial cells that are candidates to generate currents like those observed in MUC2-negative cells are K_Ca_3.1 or Kv7.1-KCNE3 (24, 27, 43, 44). In fact, in 2 of the 16 recorded non-fluorescent cells, the voltage-independent current yielded E_rev_~ −63 mV likely reflecting the primary contribution of only a K^+^ selective conductance. These two cells were located at the bottom of their respective crypts and their current showed properties similar to K_Ca_3.1 (45). Although further pharmacological studies are required to identify the conductances contributing to cation currents in MUC2-negative cells, such currents have no features consistent with BK (Fig. 3E-F). Even if a few BK channels were present, they should be readily detected under our recording conditions due to BK channel’s large and unique unitary conductance. Unlike BK current in GCs, cation currents in MUC2-negative cells exhibited a significant rundown over a few minutes. Furthermore, MUC2-negative cells have an overall current density considerably smaller than fluorescence-identified GCs (Fig. 3G) despite having similar capacitance (Fig. 3H). The fact that none of the MUC2-negative cells (n = 16) exhibit BK currents strongly suggests that colonic absorptive enterocytes not only lack LRRC26, but also lack the BKα pore-forming subunit. The finding that no BK current was observed in mouse distal colon enterocytes was surprising and contrasts with the general assumption that BK channels are widely expressed in colonic epithelial cells. Our results are in agreement with a study in human intact crypts isolated from patient biopsies, which found BK only in a subset of non-identified human crypt cells (27).

**Figure 3.**
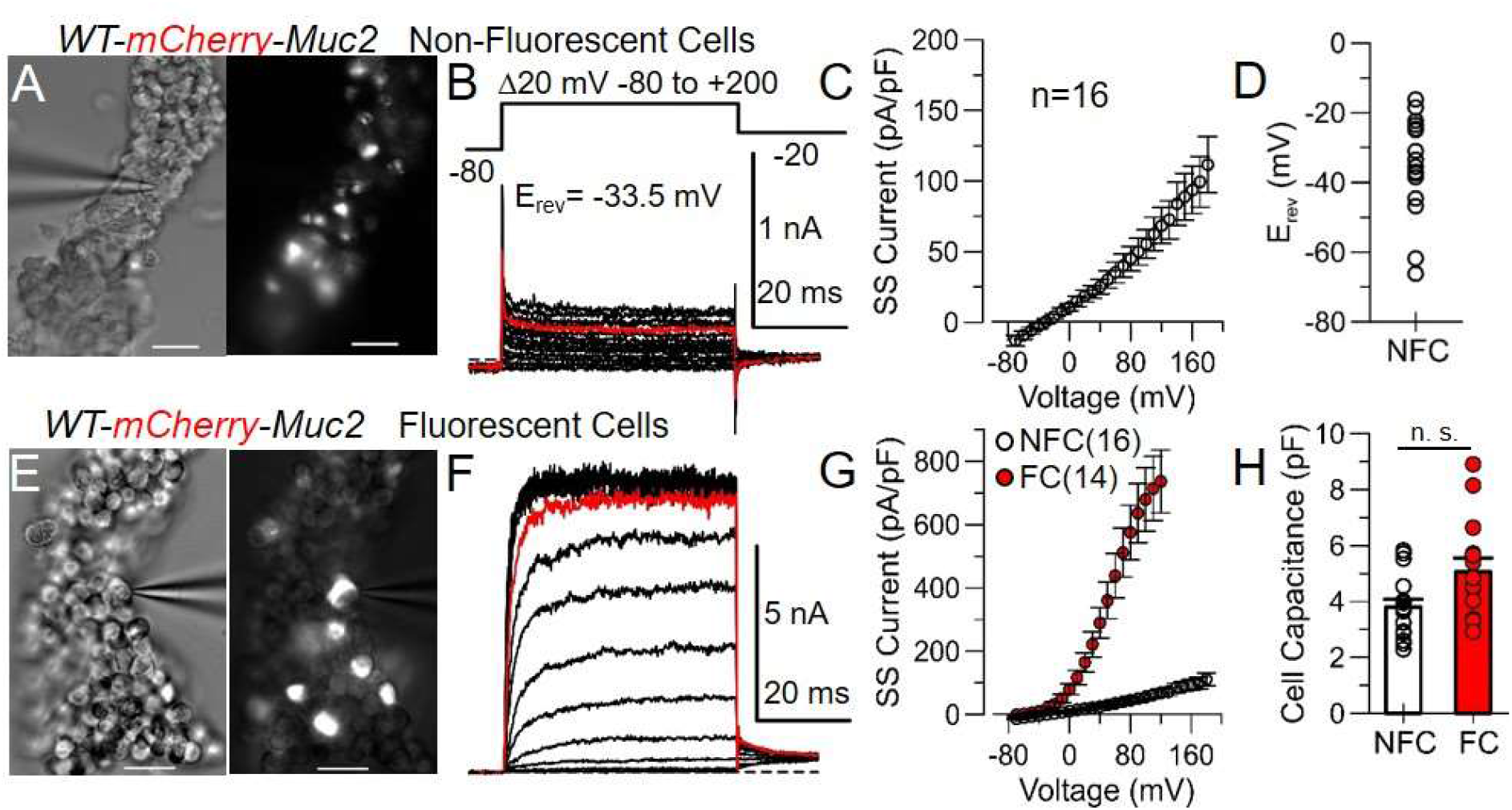
Colonic epithelial MUC2-negative cells tested lack BK channel currents. **A**, Bright field and fluorescent images of colonic crypt isolated from *WT* mouse (opening of the crypt oriented downward), with the recording pipette attached to the non-fluorescent cell whose current is shown in (**B**), scale bars are 10 μm. **B,** Representative whole-cell currents evoked by the same voltage protocol (top) used in *WT* fluorescence-identified GCs. Red trace is the current evoked by depolarization up to +100 mV. **C,** Voltage dependence of the whole cell current of non-fluorescent cells after normalization by cell capacitance (means ± SEM). **D,** Reversal potential of currents from individual non-fluorescent cells. (**E-F**) Images and representative whole-cell currents of a *WT* fluorescence-identified GC, showing the differences in current waveform and magnitude when compared with non-fluorescent current. Note the considerable difference in amplitude reflected in the scales. **G-H**, Comparison between non-fluorescent cells (NFC) and fluorescent cells (FC) (means ± SEM) in terms of current densities (**G**) and cell capacitance (**H**). Values of current densities are significantly different at membrane potentials above −20 mV (P<0.001), while capacitance values are not (P=0.054), as compared by KS-test.

### LRRC26-associated BK channels contribute to transepithelial current under resting physiological conditions

BK channels are involved in potassium secretion in distal colon of several mammalian species (27, 30, 37, 46) and results above demonstrate that LRRC26-containing BK channels have activation properties that would likely allow them to contribute to K^+^ flux under physiological conditions. Under resting physiological conditions, Ussing chamber experiments have shown that active BK channels contribute to the net basal ionic current across distal colonic mucosa *ex-vivo* (30, 37). This basal current is sensitive to both block of BK channels via luminal application of pharmacological agents (30, 37) or the genetic ablation of the BK α-subunit (30). Since LRRC26 uniquely defines the functionality of GC BK channels and we did not detect BK channels in MUC2-negative cells, we hypothesized that the contribution of BK channels to the basal transepithelial current should depend on the presence of LRRC26. Thus, genetic ablation of LRRC26 should mimic the effects of BK α-subunit KO. We therefore measured short circuit current (I_SC_) across pieces of distal colon stripped of smooth muscle after equilibrating the tissues for 30 min in symmetrical Krebs solution. Similar to previously reported values for basal I_SC_ of mouse distal colonic mucosae (30, 47, 48), our *WT* tissues showed on average a basal I_SC_ = 21.6 ± 10.0 μA/cm^2^ (mean ± SD, n=10) (Fig. 4A-B, E). However, *Lrrc26^-/-^* mucosae showed a significantly more positive basal I_SC_ = 104.0 ± 39.5 μA/cm^2^, n=11 (Fig. 4A, C, E), statistically similar to I_SC_ of *BK^-/-^* tissues (basal I_SC_ = 112.6 ± 25.7 μA/cm^2^, n=6, Fig. 4A, D, E), and in agreement with previously reported basal I_SC_ of *BK^-/-^* distal colonic mucosa (30). Our results are consistent with apical LRRC26-associated BK channels underlying a basal K^+^ efflux in *WT* tissues. Since a cationic current in the serosal-mucosal direction would contribute a negative (downward) current to the net I_SC_, its absence in tissues lacking any of the subunits required for functionally normal GCs BK channels would lead to a more positive net basal I_SC_. To further test this hypothesis, we examined the effects of the pharmacological block of BK channels on the basal current by luminal application of 5 mM TEA. TEA offers several advantages in terms of accessibility and kinetics of block for this type of experiment in comparison to more specific extracellular blockers of BK channels such as Iberiotoxin (IbTx). TEA is a positively charged, highly soluble, small molecule acting as a fast blocker and, therefore, does not require long exposure times characteristic of drugs such as IbTx or paxilline. Since BK channels exhibit the highest affinity for extracellular TEA among all K^+^ channels (K_d_ ~ 1 mM) (49, 50) the use of a low [TEA] allows a fairly specific, fast, and readily visualized effect. As predicted, 5 mM TEA induces a prompt upward change in *WT* basal I_SC_ that reaches a steady state within a couple of minutes (ΔI_SC_ (*WT*)=19.4 ± 9.2 μA/cm^2^, n=10 (mean ± SD) (Fig. 4A-B,F), while producing no significant change in Isc of either *Lrrc26^-/-^* or BK^-/-^ tissues (ΔI_sc_(*Lrrc26^-/-^*)= −3.8 ± 9.2 μA/cm^2^, n=11; Δ I_SC_ (*BKα^-/-^*) = −2.3 ± 10.8 μA/cm^2^, (n=6) (Fig. 4A, C-D, F). In order to verify the viability of all the tissues assayed, the experiments ended with the application of 10 μM of luminal forskolin. The presence of a 5 mM TEA-sensitive current under resting conditions, which is absent in both KO mouse models, supports the conclusion that LRRC26-associated BK channels of GCs underlie the K^+^ secretion occurring in distal colon at resting physiological conditions.

**Figure 4.**
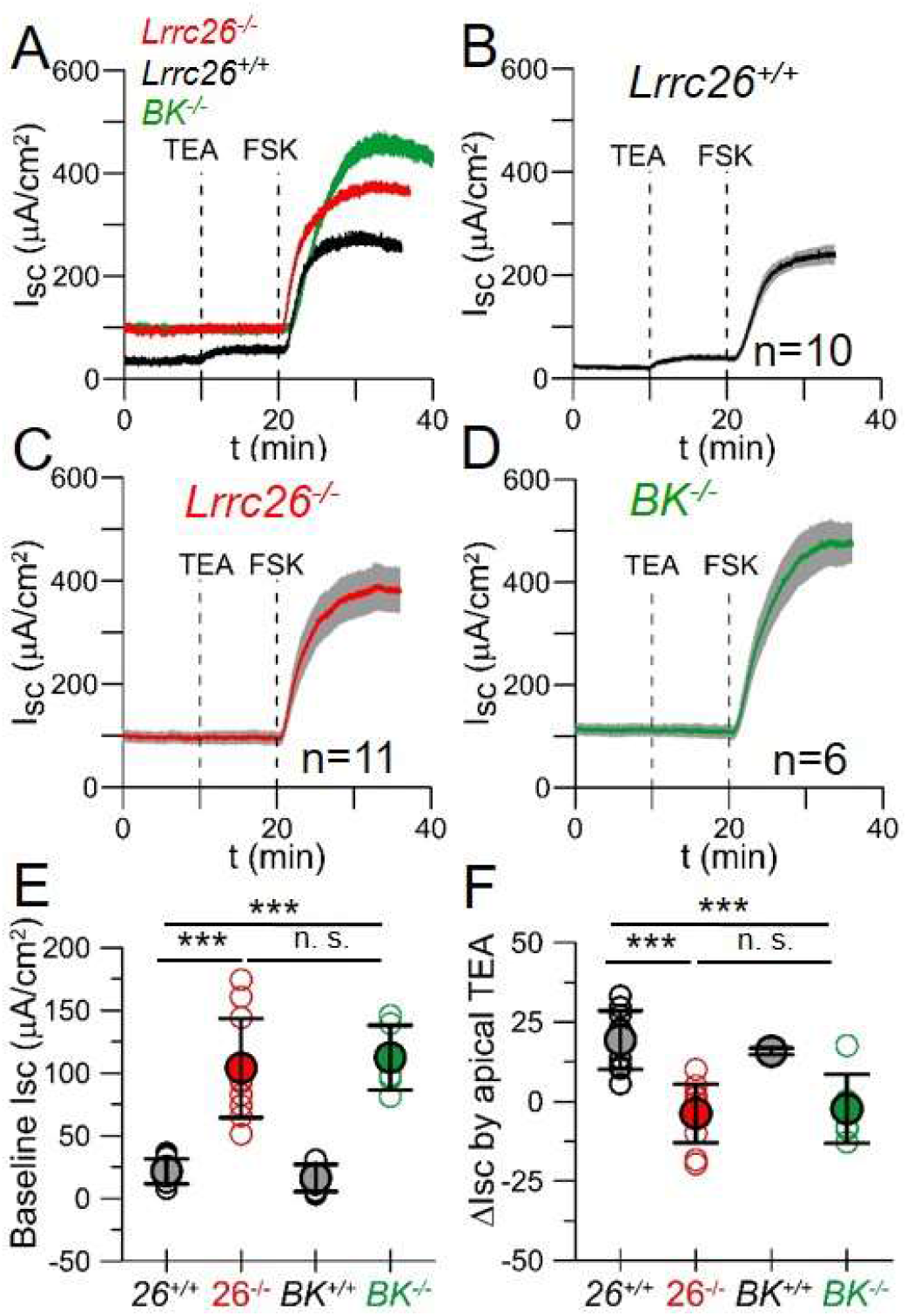
LRRC26-associated BK channels contribute to the resting K^+^ secretion in distal colonic mucosa. **A**, Representative transepithelial currents (Isc) recorded under voltage clamp at 0 mV in distal colonic mucosae from different genotypes. In all cases baseline Isc was recorded for 10 min followed by luminal application of 5 mM TEA and 10 μM forskolin at the indicated times. **B-D**, Averaged Isc (line) ± SEM (gray shadow) obtained from independent experiments for *Lrrc26^-/-^* (**B**), *Lrrc26^+/+^* (**C**), and *BK^-/-^* (**D**). **E,** Baseline Isc (mean ±SD) in the different genotypes. Sets of values were compared with Anova with Tukey’s correction. For *Lrrc26^+/+^* vs. *Lrrc26^-/-^*, P<0.0001 (n=10 vs. n=11); for *Lrrc26^+/+^* vs. *BK^-/-^*, P<0.0001 (n=10 vs. n=6); for *Lrrc26^-/-^* vs. *BK^-/-^*, P= 0.931 (n=11 vs. n=6). **F,** Response to luminal TEA (mean ±SD). For *Lrrc26^+/+^* vs. *Lrrc26^-/-^* P<0.0001 (n=10 vs. n=11); for *Lrrc26^+/+^* vs. *BK^-/-^*, P<0.0001 (n=10 vs. n=6); for *Lrrc26^-/-^* vs. *BK^-/-^*, P= 0.9878 (n=11 vs. n=6).

### The absence of LRRC26 increases susceptibility to DSS-induced colitis

Given the key role of LRRC26 in defining the functional properties of GC BK channels, we sought to investigate whether LRRC26-associated BK channel activity in GCs has implications for colonic epithelial homeostasis. Although *Lrrc26^-/-^* mice exhibited no differences from WT mice in terms of gross appearance, colon histology (including crypt morphology and number of GCs) (Fig. 1E-F, Fig. S3), or fecal microbiota (Fig. S4), we compared the responses of *Lrrc26^-/-^* and *WT* littermates in an experimental model of induced colitis. Since our data indicate that the number of *Lrrc26*-positive GCs display a gradient increasing towards the distal colon, we chose a colitis model that involves a disruption of the epithelial barrier with inflammation mainly localized to the distal colon, as is characteristic of the acute colitis induced by DSS (51, 52). Sets of littermates were provided with drinking water containing 2.5% DSS for 7 days, and then returned to regular water. Starting on the 5^th^ day of treatment both *Lrrc26^-/-^* and *WT* mice displayed a typical drop of body weight (51), accompanied by increasingly loose stools with bloody diarrhea becoming apparent around day 7. While in most *WT* mice the diarrhea ceased within about 24 h after switching to regular water with weight beginning to recover on day 10, in *Lrrc26^-/-^* mice bloody diarrhea and body weight loss persisted. Between days 9-10 a significant fraction of *Lrrc26^-/-^* mice began to show clinical signs of morbidity, including a hunched posture and inactivity to the point where they died or required euthanasia due to a weight loss over 30% of the initial value. By day 13 around 50% of *Lrrc26^-/-^* treated with 2.5% DSS died or required euthanasia in comparison to <10% of *WTs* (Fig. 5A-B). Increasing the DSS concentration to 3.5% DSS resulted in 100 % lethality of *Lrrc26^-/-^* versus <30% in *WT* littermates (Fig. 5C). We also compared BK^-/-^ and *WT* littermates using 2.5% DSS; 100% of BK^-/-^ mice died or required euthanasia vs. none of *WT* mice (Fig. S5). A more severe phenotype for BK^-/-^ mice is not unexpected, given the strong expression of BK α-subunit in non-epithelial colonic cells including the smooth muscle cell layer (30) and Cajal’s cells in the *lamina propria* (53).

**Figure 5.**
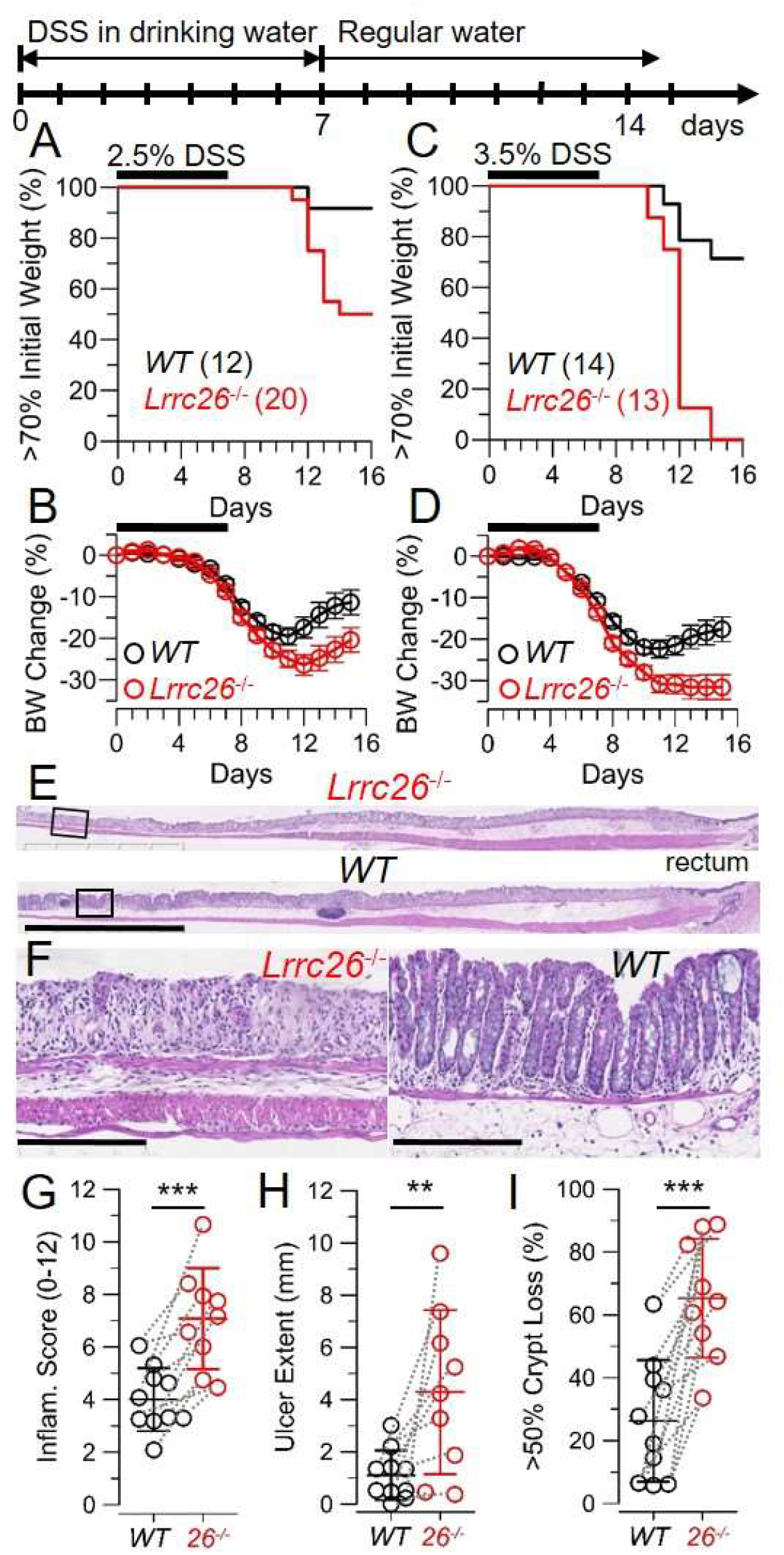
Genetic ablation of LRRC26 dramatically enhances susceptibility to DSS-induced colitis. **A-D**, Survival plots and change in body weight curves of mice treated for 7 days with 2.5% (**A-B**) or 3.5% DSS (MP Biomedicals, MW: 36-50 KDa) (**C-D**). In accordance with guidance from WUSTL-animal studies committee, animals whose body weight dropped > 30% of their initial value were considered terminal and euthanized. Graphs compile results from 4 independent assays where 8 litters (with both *Lrrc26^-/-^* and *WT* littermates) were treated with 2.5% DSS or 2 assays where 4 litters were treated with 3.5%. Mice were 10-14 weeks old at the time of the treatment and both sexes were included. In body weight plots (mean ± SEM), for mice that died or required euthanasia, weight at the time of death was considered the same until the end of the experiment. Curves were compared using a two-way ANOVA that yielded P=0.0117 for 2.5% and P=0.0073 for 3.5% DSS. **E**, Representative images of H&E stained most distal segment of colon from *Lrrc26^-^* and *WT* (littermates) at the 7^th^ day of treatment (bars: 2.5 mm). **F**, Areas within the squares in (E), (bars 250 μm). **G-I**, Histophatologic colitis severity in the distal colon of DSS-treated mice scored from sections as in E-F. Inflammation score (**G**), total ulcer extention (**H**) and fraction of epithelial surface with 50% or more of crypt loss (**I**), represented as mean±SD. Dotted lines connect results from siblings. Sets of values were compared using two-tailed t-test that yielded P=0.00054, P=0.0071, and P=00036 for comparisons in G, H, and I, respectively.

The differences in colitis severity of *Lrrc26^-/-^* vs. *WT* mice treated with DSS were confirmed by histology (Fig. 5E-H). Hematoxylin-Eosin (H&E) stained sections showed that, at day 7 of 2.5% DSS treatment, *Lrrc26^-/-^* mice already have a severe crypt depletion in the distal colon in comparison with *WT* littermates (Fig. 5E-F, I). In addition to the crypt loss, *Lrrc26^-/-^* mice show increased inflammation (Fig. 5E-F, H), more extensive ulcerations (Fig. 5E-F, H) and larger infiltration of immune cells from the *lamina propria* (Fig. 5E-F). Overall, these results demonstrate that *Lrrc26^-/-^* mice exhibit increased colitis severity and suggest a protective role for LRRC26-BK channels in the colonic epithelium.

## Discussion

The present study establishes the following: 1) In colonic GCs, BK channels are associated with LRRC26 regulatory subunits and represent the major cation current of these cells; 2) the shifted gating properties of LRRC26-containing BK channels allow them to be active at expected normal GC resting conditions, which was also confirmed by the impact of LRRC26 KO on transepithelial ion flux; 3) colonic absorptive enterocytes within crypts lack BK channel currents; 4) the absence of LRRC26 in GC BK channels results in a severe DSS-induced colitis phenotype, also occurring with complete KO of the BKα subunit. The DSS-induced colitis is associated with a significantly enhanced loss of crypts and GCs in the distal colon of *Lrrc26^-/-^* in comparison to *WT* littermates.

### LRRC26 is a critical component of GC BK channels which underlie the major cation current in these cells

Most existing models that summarize the interplay among different ion transport mechanisms in intestinal epithelial cells are focused on absorptive enterocytes (8, 46), whereas ion transport in GCs has been largely overlooked. Yet, there are several lines of evidence supporting the presence of BK channels in colonic epithelial cells of several species including mice (30), rats (22, 43), guinea pigs (37) and humans (24, 27, 54), although without identifying the cell type. In rodents, colonic epithelial BK channels play a prominent role in K^+^ secretion under both basal conditions (30) and upon stimulation with purines (30), adrenaline (28), aldosterone (29, 55) or in response to high dietary K^+^ load (22, 29). However, a long lasting unanswered question has been how these epithelial BK channels can be activated under physiological conditions. BK channels composed exclusively of the pore-forming BKα subunit would require both elevation of cytosolic Ca^2+^ and membrane depolarization in order to achieve activation necessary for its physiological roles (56). Therefore, in order for BK channels to be active under physiological conditions of colonic epithelial cells (reported to be ~120 nM intracellular Ca^2+^ at resting (57) and ~500 nM under muscarinic stimulation (58) in GCs), it seems likely that a regulatory subunit may be required. In this study we demonstrate that the regulatory subunit LRRC26 is a critical component of BK channels in colonic GCs, which allow them to contribute to epithelial K^+^ efflux even at resting conditions, although cytosolic elevations of Ca^2+^ triggered by secretagogues or other stimuli would further enhance GC BK-mediated K^+^ fluxes. Another surprising and unexpected finding of this study concerns the cell type location of colonic epithelial BK channels. It has been suggested that epithelial BK channels fulfill their above mentioned role in K^+^ secretion by widespread expression in colonic epithelial cells including enterocytes (8, 46). Here, we provide evidence that unambiguously indicates that, at least in mouse distal colonic epithelium, cells belonging to the GC lineage contain BK current, while BK current is absent in all MUC2-negative cells tested. Since enterocytes are the major population of cells negative for MUC2 in distal colon, the conditions of our experiments should have readily revealed BK current in enterocytes, if it were present. In agreement with our results, previous evidence suggests that location of BK channels in GCs likely also occurs in human distal colon epithelium (27). Specifically, BK and K_Ca_3.1 currents are the major K^+^ currents in different subsets of cells from human intact crypts (27). Although those cell types were not identified, only K_Ca_3.1-cells developed a CFTR-mediated response to increased levels of intracellular cAMP. The absence of a CFTR response in human crypt cells having BK currents suggests that CFTR, a protein well known to be expressed in enterocytes, and BK channels are located in different types of epithelial cells in human distal colon (27). Our evidence regarding the segregation of BK channels between distinct epithelial cell types, besides challenging previous general expectations about BK cellular localization in colonic epithelium, suggests that the functional contributions of BK channels are tightly linked to the functional roles of GCs.

### What might be the role of BK current in GC function?

The segregation of BK channels among colonic epithelial cells shares an interesting parallel with the previously observed probable segregation of two Ca^2+^-dependent anion conductances, mediated by BEST2 and TMEM16A (also known as ANO1) in mouse colonic epithelium (59). In this work, Ca^2+^-activated anion currents with properties similar to heterologously expressed BEST2 currents were the major anion current in a subset of dissociated epithelial cells presumed to be GCs. In different epithelial cells, presumed to be enterocytes, the principal Ca^2+^-activated anion current exhibited properties similar to heterologously expressed TMEM16A currents. BEST2, which has a high bicarbonate permeability relative to other anions, seems to be involved in the HCO_3_^-^ secretion upon cholinergic stimulation (59). Besides both being specifically expressed in murine GCs, BK channels and BEST2 share a number of other features. First, both are activated by elevations of cytosolic Ca^2+^; second, *Best2* shares with *Lrrc26* a similar increased proximal to distal gradient of expression; third, *Best2^-/-^* mice also exhibit a DSS-colitis phenotype although less severe than that occurring in *Lrrc26^-/-^* mice. One difference is that, although LRRC26-containing BK channels contribute to basal K^+^ flux as revealed by the BK-dependent component of basal I_SC_, BEST2 probably exhibits little activation until intracellular Ca^2+^ is elevated. Yet, it seems reasonable to think that the coincident activation of K^+^ flux through BK channels and bicarbonate fluxes through BEST2 channels are somehow critical in supporting mucus secretion and the extracellular conditions of Ca^2+^ and elevated pH critical to mucus expansion.

In addition to BEST2, GC BK channels might act in concert with other ion transport entities proposed to be implicated in colonic mucus secretion. For example, activation of luminal purinergic G-protein-coupled (P2Y) receptors by ATP or UTP in mouse colon stimulates a K^+^ secretion mediated by BK channels (30, 60). Luminal ATP stimulation also induces mucus secretion by intestinal GCs (57), which is compromised in mice with KO of TMEM16A specifically in intestinal epithelial cells. In fact, this mouse model exhibits an accumulation of mucus in intestinal GCs (57). An increase in BK channel-mediated K^+^ flux during ATP-mediated mucus secretion seems plausible since the activation of P2Y receptor produces an elevation in cytosolic Ca^2+^ concentration measured in colonic GCs (57, 58). Similar mucus accumulation in GCs has also been observed in a mouse model that recapitulates a human mutation in NKCC1 (21). Both, the mouse model and patients carrying this NKCC1 truncation mutation also exhibit deficient exocytosis of mucus granules from GCs, mucus attached to epithelium, and a thinner colonic mucus layer (21). Although it is not clear yet whether TMEM16A and/or NKCC1 influence mucus secretion by intrinsic expression in GCs or by a paracrine effect from the neighboring enterocytes, if the first case is true, K^+^ flux through BK channels might act in concert with the anion transport generated by those entities.

LRRC26-containing BK channels may act in concert, acting as a necessary counterion, of an anion conductance that perhaps is directly involved in the initiation of mucus exocytosis, or is critical for appropriate mucus maturation. *Lrrc26^-/-^* untreated mice do not show evidence of mucus accumulation, which may suggest a role downstream of exocytosis. Remaining important unanswered questions include: 1) Is there a change in mucus properties and-or mucus secretion in the *Lrrc26^-/-^* mice? 2) Is there any residual non-BK K^+^ current in GCs? 3) What are the K^+^ fluxes in enterocytes? Answers to these questions will allow a better understanding of the role of ion transport in colonic mucus secretion.

### LRRC26-associated BK channels have a protective effect in an experimental model of DSS-induced colitis in mouse and perhaps in human IBD

Our experiments in *Lrrc26^-/-^* mice indicate that the activity of LRRC26-associated BK channels has a protective effect against colitis induced by DSS. Not surprisingly, *BK^-/-^* mice exhibit a more profound colitis phenotype with the same treatment (Fig. S5, Fig. 5). These differences may be explained by the high levels of expression of BKα subunit in smooth muscle cells (30, 61), and interstitial cell of Cajal (53), which suggest that *BK^-/-^* mice might also have a compromised intestinal motility.

Given the strong phenotype observed as result of genetic ablation of the subunits contributing to colonic epithelial BK channels, what are the implications of epithelial BK channels in human IBD? In Ulcerative Colitis (UC) patients, expression of BK α subunit in colonic crypts is enhanced (54). Whether the upregulation is an initial step in pathology, is part of a compensatory response, or involves only some or all epithelial cells are all unknown. At present, nothing is known about LRRC26 expression in UC. This is important information considering that colonic epithelial BK channels lacking LRRC26 are likely non-functional under physiological conditions. One might imagine that some variables such as diet, aging, or stress-related factors could impact on LRRC26 expression, and potentially influence the susceptibility to UC. Intriguingly, work with LRRC26-containing BK channels expressed heterologously indicate that LRRC26 and BKα subunit interactions can be labile (42). Therefore, we addressed the possibility that *LRRC26* gene expression may change in different UC disease states. Analysis of a public database containing gene expression profiles from colon of patients with IBD (62) reveals a significantly reduced expression of *LRRC26* in samples of patients with active UC in comparison to patients with inactive disease (Fig. S6). As expected for a GC specific protein, *LRRC26* expression positively correlates with *MUC2* expression among all samples analyzed. Consistent with having a protective role, *LRRC26* expression negatively correlates with the expression of the inflammatory cytokine IL-6 in samples of patients with active disease. These results suggest that understanding the protective role of LRRC26-associated BK channels against colitis may provide a new opportunity to understand IBD pathophysiology.

## Materials and Methods

### Animal care

Animals were handled and housed according to the National Institutes of Health Committee on Laboratory Animal Resources guidelines. All experimental protocols were approved by the Institutional Animal Care and Use Committees of Washington University in St Louis (protocol #20180288). All mice were on a C57BL/6J background. The *Lrrc26^-/-^* mouse strain is a global KO previously described (32) and is available from the KOMP Repository (WWW.KOMP.org) with the neomycin-resistance selection cassette deleted. The mouse referred as *BK^-/-^* is the *Kcnma1^-/-^* mouse line, a kind gift of Dr. Andrea Meredith (University of Maryland, Baltimore, MD). The mCherry-Muc2 mouse line was kindly shared by Dr. Gunnar C. Hansson (University of Gothenburg, Sweden).

### X-Gal staining

Fresh entire colons (from anus to caecum) from *Lrrc26^-/-^* and *WT* mice were dissected and rinsed with cold 0.1M phosphate buffer, pH=7.4 (PB). Then, colons were opened longitudinally along the mesenteric line and quickly immersion-fixed, by pinning flat with the mucosal surface facing up, in a chamber containing fixative solution, at 4°C. The fixatives procedures were 0.2% glutaraldehyde for 24 h or 4% paraformaldehyde for 1 h, both dissolved in PB. Fixed tissues were transferred to ice-cold 30% sucrose in PB and kept at 4°C for 48 hours. Colonic pieces of 3×18 mm were mounted in OCT block and frozen in dry ice. 20 μm frozen tissue sections were then prepared in a cryostat at −20°C. Slides were washed once briefly with 0.1 M PB and then once with 37°C X-Gal dilution buffer (5 mM Potassium Ferricyanide, 5 mM Potassium Ferrocyanide, 2 mM Magnesium Chloride, 0.1% Tween-20 in PBS (pH 7.4)). Just before staining, X-Gal stock solution (4% (w/v) Bluo-Gal (Invitrogen, 15519-028) in dimethylformamide) was diluted at 1:40 in 37°C X-Gal dilution buffer to prepare the fresh X-Gal working solution. Tissue sections were then incubated in the X-Gal working solution at 37°C in a humidity box. Incubation times of 1, 2, 3, 6, 18 and 24 h were assayed. Using glutaraldehyde fixation, 1-2h is enough to develop a strong and specific signal while 3h or longer incubation times result in unspecific signal in *WT* tissues. Paraformaldehyde fixed tissues requires 18-24 h to develop a signal. Finally, sections were rinsed in deionized water, and some sections were counterstained with Periodic Acid-Schiff (PAS) Staining System (395B-1KT; Sigma-Aldrich) before being mounted with aqueous mounting medium.

For quantification of the number of stained cells along colon, from distal end until the beginning of the ridges in the proximal colon, blue stained cells per crypt were counted in 5 consecutive well oriented crypts, at intervals of 5 mm.

### Morphological and histological analysis

Fresh entire colons (from anus to caecum) from *Lrrc26^-/-^* and *WT* littermates were dissected, rinsed with cold PBS, opened longitudinally along the mesenteric line and pinned flat in a chamber containing 4% paraformaldehyde in PBS. Tissues were fixed overnight at 4 °C then transferred to 70 % ethanol before processing for paraffin embedding. 5 micrometers serial sections were prepared and stained with H&E or PAS. Slide imaging was performed in a Hamamatsu Nanozoomer 2.0 HT System hosted by Washington University Hope Center. Morphological measurements were performed as has been previously described (63), crypt depth was quantified as the average from at least 25 well oriented crypts in the distal colon per animal; abundance of GCs was measured from PAS stained tissues, and was calculated from at least 25 well oriented crypts in the distal colon, as the percentage of GCs (stained in dark pink) regarding the total number of cells of a crypt (counting the nuclei). Colitis severity in DSS-treated mice was scored as have been described previously (63, 64).

### Preparation of crypts for whole-cell recording

Intact crypts from distal colon were isolated by a modification of a Ca^2+^ chelation protocol previously reported (27, 65). Mice of 10-16 week-old, both sexes, were used, one mouse per day of recording. Colon was quickly dissected, rinsed with cold PBS to remove fecal content and opened longitudinally following the mesenteric line. The most distal 2 cm were cut, placed in a small glass flask containing 10 mL of Ca^2+^-chelating solution and incubated at 37 °C. Ca^2+^-chelating solution was always used freshly made and contained (in mM): 20 HEPES, 10 EDTA, 112 NaCl, 5 KCl, 3 mM dithiothreitol and adjusted to pH=7.1 with TRIZMA base. After 3 minutes of incubation at 37°C, the piece of colon was briefly and gently stirred within the solution, and further incubated at 37°C for another 3 minutes. Then, after changing the piece of tissue to a new small flask containing fresh Ca^2+^-chelating solution pre-incubated at 37°C, the suspension was shaken vigorously for 3-4 sec using a spatula, repeatedly 5-6 times, to liberate intact into the solution. Crypts were collected by centrifugation at 150g for 3 min at 4°C and washed twice with cold Crypt-storage solution containing (in mM): 100 K-gluconate, 30 KCl, 20 NaCl, 1.25 CaCl_2_, 1 MgCl_2_, 10 HEPES, 5 Glucose, 5 Na-Pyruvate, 5 Na-Butyrate, supplemented with 1g-L BSA, pH=7.4 adjusted with KOH. The pellet resulting from the last centrifugation was resuspended in 0.5 mL of Crypt-storage solution. 0.1 mL of the final suspension containing the crypts was seeded in coverslips (diameter 12 mm) and allowed to attach for 1h at RT before beginning the patch clamp recordings. The coverslips had been previously treated with a dilution 1/10 of Matrigel (Corning, cat #354234) in PBS for at least 30 min at 37 °C and rinsed with PBS just before seeding the crypt suspension. All experiments were obtained within 1-5 hours after crypt dissociation.

### Patch clamp recordings

Standard whole-cell recording methods were done using a Multiclamp 700B amplifier (Molecular Device, San Jose, CA USA). Data acquisition was performed using a 16 bit analogue-digital converter and voltage stimulation protocols were accomplished by using Clampex 9.0 (Molecular Devices) with analysis of waveforms done via Clampfit. Patch-clamp pipettes were made from borosilicate glass and coated with Sylgard. Typical pipette resistances after heat-polishing typically were of 3 MΩ. Following whole-cell access, cells were used if the series resistance (R_s_) was less than 10 G Ω. R_s_ was 90 % compensated. Current records were filtered at 10 kHz. Recording solutions contained physiological gradients of Na^+^ and K^+^, while most of Cl^-^ was replaced by glutamate to minimize the contribution of any Cl^-^ current. The bath solution (extracellular) consisted of (in mM): 135 Na-Glutamate, 5 K-Glutamate, 2 CaCl_2_, 2 MgCl_2_, 10 HEPES (pH = 7.2). The pipette solution (intracellular) consisted of (in mM): 135 mM K-glutamate, 10 HEPES and, 5 mM EGTA + 3 mM Ca^2+^ which results in a free Ca^2+^ solution of 250 nM (pH=7.2). For these nominal ionic gradients, theoretical E_K_ = −84.5 mV. All the experiments were at nominal room temperature (22-25 °C). Tetraethylammonium (*Sigma*) or paxilline (*Tocris Bioscience*, Bristol, UK) were added (from concentrated stock solutions freshly prepared) to the extracellular solutions at final concentrations given in the text.

G-V curves of fluorescence-identified GCs were generated from the tail currents measured following a repolarization step at −20 mV for *WT* cells or at +20 mV for *Lrrc26^-/-^* cells. Tail current amplitude at 400 μs following the nominal repolarization step to minimize contribution of uncompensated signals. G-V curves were fitted to a single Boltzmann distribution:

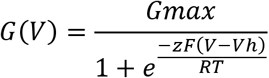

where V_h_ represents the voltage of half activation and z is the valence of the voltage dependence. F and R are the Faraday’s and Gas’s constants, respectively and T is the temperature.

### Ussing chamber experiments

We used the most distal 2 cm of the mouse colon. After colon dissection, smooth muscle layer was removed by stripping, and a piece of mucosa was mounted in an Easy Mount Ussing chamber (Physiologic Instruments Inc.) using an insert with an aperture of 0.1 cm^2^ (P2303A, Physiologic Instruments). The standard Krebs solution contained 119 mM NaCl, 2.7 mM KCl, 23 mM NaHCO_3_, 1.25 mM Na_2_HPO_4_, 1.8 mM CaCl_2_, 1.5 mM MgSO_4_, 0.5 mM ascorbic acid, and 10 mM glucose. The reservoirs were continuously gassed with 5% CO_2_ and 95% O_2_ and maintained at 37°C by water jackets. Isc was measured using an automatic voltage clamping device (EVC-4000; Physiologic Instruments Inc.) that compensates for resistance of the solution between the potential measuring electrodes. The transepithelial current was applied across the tissue via a pair of Ag/AgCl electrodes that were kept in contact with the mucosal and serosal bathing solution using 3M KCl-agar bridges. All experiments were done under voltage clamp at 0 mV. The Isc is negative when positive current flows from serosa to mucosa. The tissue was placed in the apparatus and equilibrated for 30 minutes to stabilize Isc before starting the experiment. The baseline value of electrical parameters was determined as the mean over the 10 minutes immediately prior to drug administration. A positive Isc corresponds to the net electrogenic secretion of anions or the net electrogenic absorption of cations. Forskolin (10 μM) were used on the luminal side for confirming the viability of the tissues.

### DSS-induced colitis assay

Sets of siblings (both sexes included), 10-14 weeks old were treated for 7 days with drinking water containing 2.5% or 3.5 % Dextran Sulfate Sodium (DSS) (MW:36-50 kDa, MP Biomedicals), then switched to regular drinking water. Body weight and diarrhea score were monitored daily. Following guidelines Washington University DCM committee, mice with body weight <70% of the initial weight was considered terminal and euthanized. At day 7 of DSS treatment some pairs of siblings were sacrificed for histological comparison. Colitis severity was scored in the distal third of colons as has been described previously (63, 64), following a score (0-12) based on ulcer extent (0-4) + % crypt loss (0-4) + degree of edema (0-4). The total ulcer extension (mm) and the percentage of epithelial surface with >50% of crypt loss were plotted independently.

### Statistical Analysis

For comparisons between two distributions, a two-tailed unpaired Student’s t-test or a Kolmogorov-Smirnov (KS) test was employed. When multiple comparisons were required, an ANOVA with Tukey’s correction was used. For comparison of body weight curves, a two-way ANOVA with Geisser-Greenhouse correction was used. The Geisser-Greenhouse correction was required due the lack of sphericity in the data caused by the 30% limit in the body weight drop measurements.

## Author contributions

V.G.-P., M.A.C. and C.J.L. designed research; V.G.-P, P.L.M-E, M.S.-R. and N.B. performed research. X.-M.X, H.H. and J.K.G. contributed reagents/analytical tools. V.G.-P, D.A, A.C.C. analyzed data; V.G-P and C.J.L wrote the paper. All authors revised and edited the manuscript.

## Acknowledgements

This work was supported by the Lawrence C. Pakula, MD IBD Education & Innovation Fund IA-2018-10-IBD-2 to VGP and grant GM-081748 to CJL. *Kcnma1* KO mice were kindly provided by Dr. Andrea Meredith, Univ. Maryland-Baltimore. We acknowledge the Swedish Research Council grant 2014-00366 to JKG and grant 2017-00958 for the generation of the mCherry-Muc2 transgenic mouse, which was kindly provided by Dr. Gunnar C. Hansson, Univ. of Gothenburg, Sweden. We thank Dr. Thaddeus S. Stappenbeck for useful discussions. Histology services were provided by the Washington Univ. Digestive Research Core Center supported by grant P30DK052574. Slide imaging was performed in a Hamamatsu Nanozoomer 2.0 hosted by Washington Univ. Hope Center and supported by NIH shared Instrumentation Grant S10RR0227552. Microbiota analysis was performed at the Washington Univ. Genome Technology Access Center, which is partially supported by NCI Cancer Center Support Grant #P30 CA91842 and by ICTS/CTSA Grant# UL1TR000448 from the NCRR.

---

**Figure S1.**
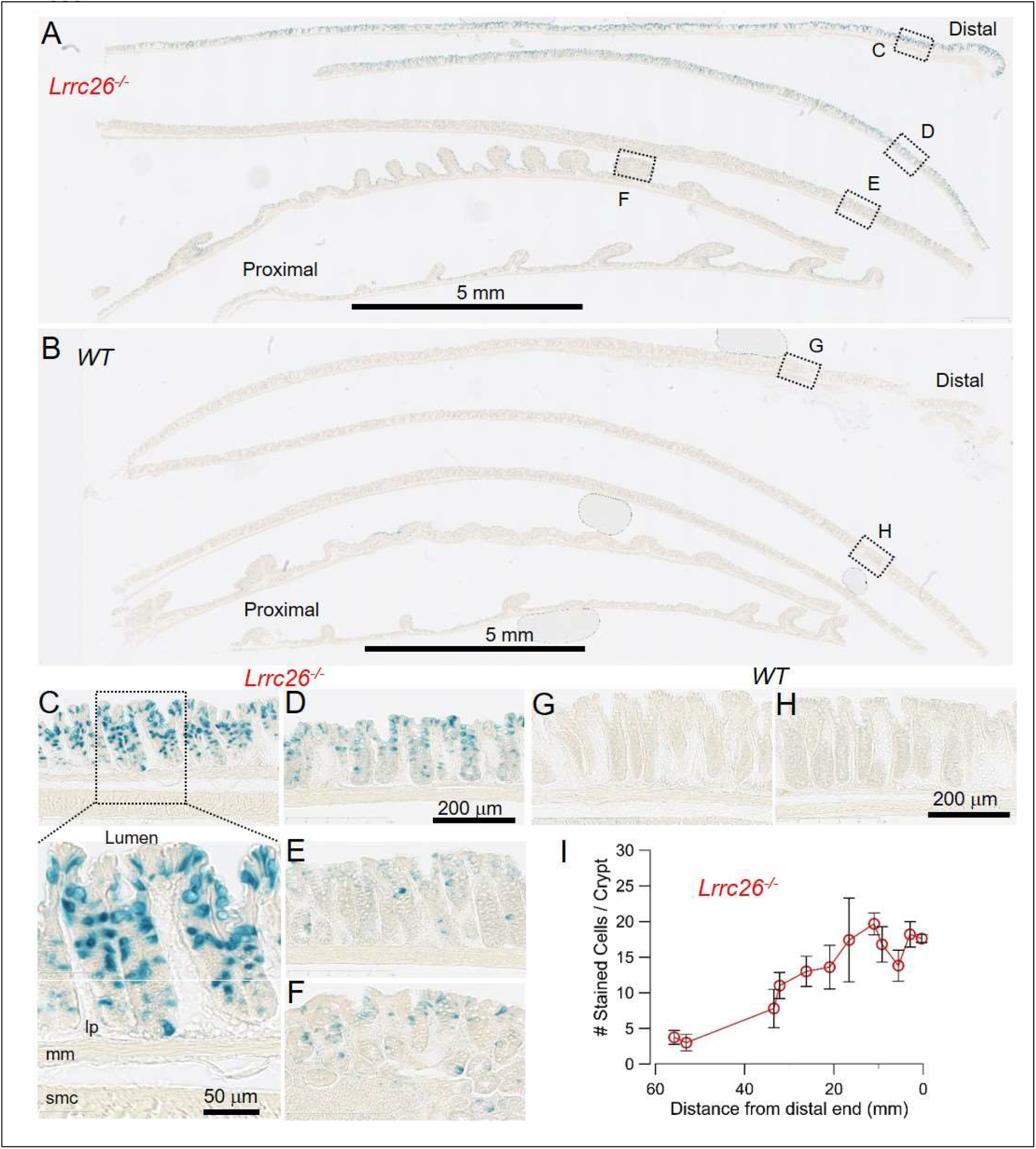
*Lrrc26* expression gradually increases in the direction proximal-distal with the highest expression in the 2/3 most distal segments. **A-B**, Images of fixed frozen sections of *Lrrc26^-/-^* (A) and *WT* (B) colons from littermates stained side-by-side with X-Gal substrate for 2 h. The entire colon (from anus to caecum), mildly fixed with 0.8% glutaraldehyde for 24 h, was divided into 5 pieces of ~1.8 cm and stacked within the OCT block with the most proximal piece at the bottom and the most distal at the top. The distal end of each piece is on the right. Panels A and B display longitudinal sections containing all 5 pieces showing a gradient of increasing β-Gal expression in the proximal-distal direction in *Lrrc26^-/-^*, while no signal was observed in *WT* tissue. Higher magnification of areas within dotted squares in *Lrrc26^-/-^* (**C-F**) and *WT* (**G-H**) pieces. In the area at the highest magnification, lumen, *muscularis mucosae* (mm), *lamina propria* (lp) and smooth muscle cell layer (smc) are indicated. **I,** Number of β-Gal positive cells as function of the distance from the anus. Each point represents the average from 5 consecutive well oriented crypts sampled at intervals of 2-5 mm (mean ±SD).

**Figure S2.**
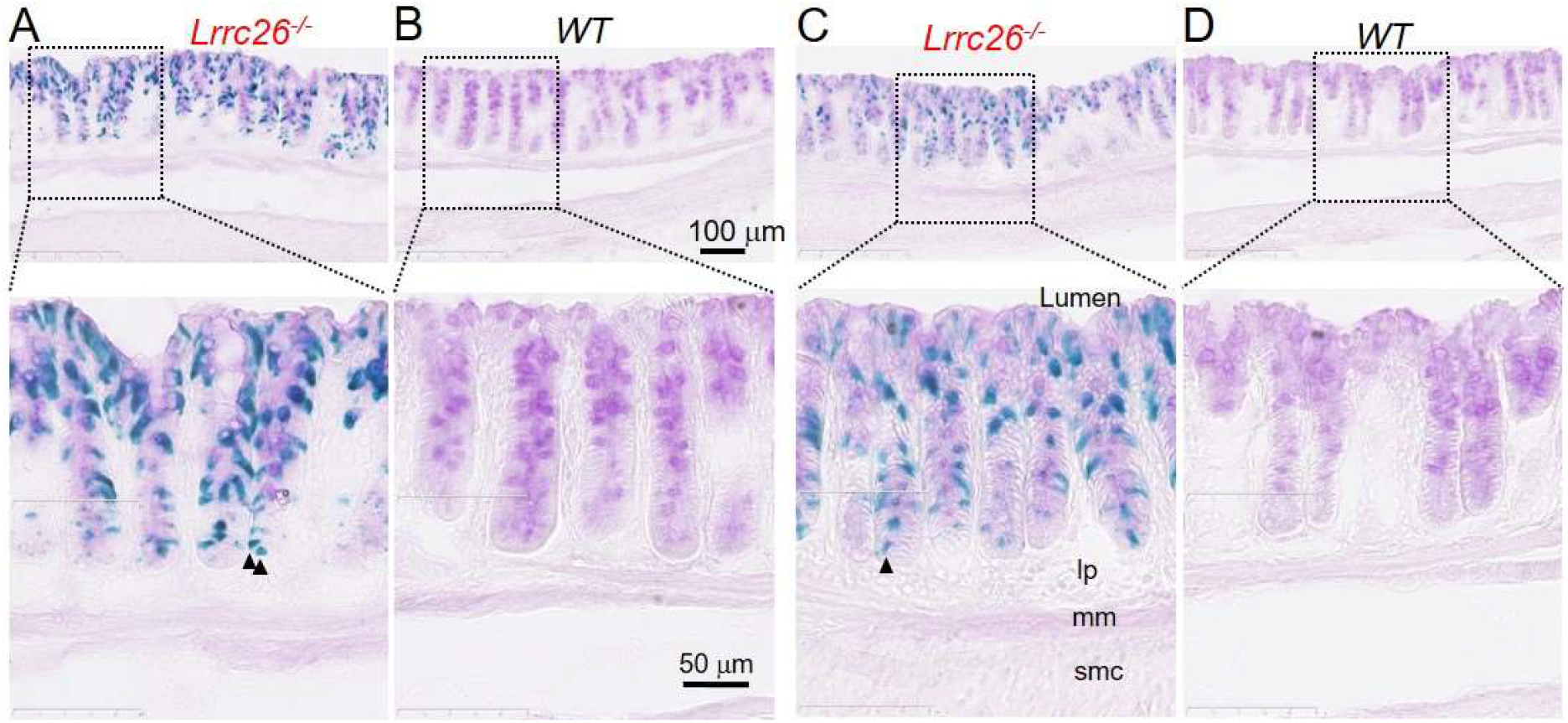
*Lrrc26* promoter activity is detected only in epithelial GCs. Representative images of PFA-fixed 20 micron frozen sections, from *Lrrc26^-/-^* and *WT* littermates treated side-by-side, stained sequentially with X-Gal for 24 h followed by PAS. Sections were taken from tissues previously fixed with 4% paraformaldehyde for 1 h at 4°C. Micrograph represent areas at 3.5 mm (**A-B**) or 10 mm (**C-D**) from the anorectal junction. Panels below are higher magnification of areas within dotted rectangles showing that X-Gal staining is specific for PAS positive cells along the crypts and is absent in *muscularis mucosae* (mm), *lamina propria* (lp) and smooth muscle cell layer (smc). Filled black triangles identify some cells at the base of crypts showing blue staining, but not appreciable PAS staining.

**Figure S3.**
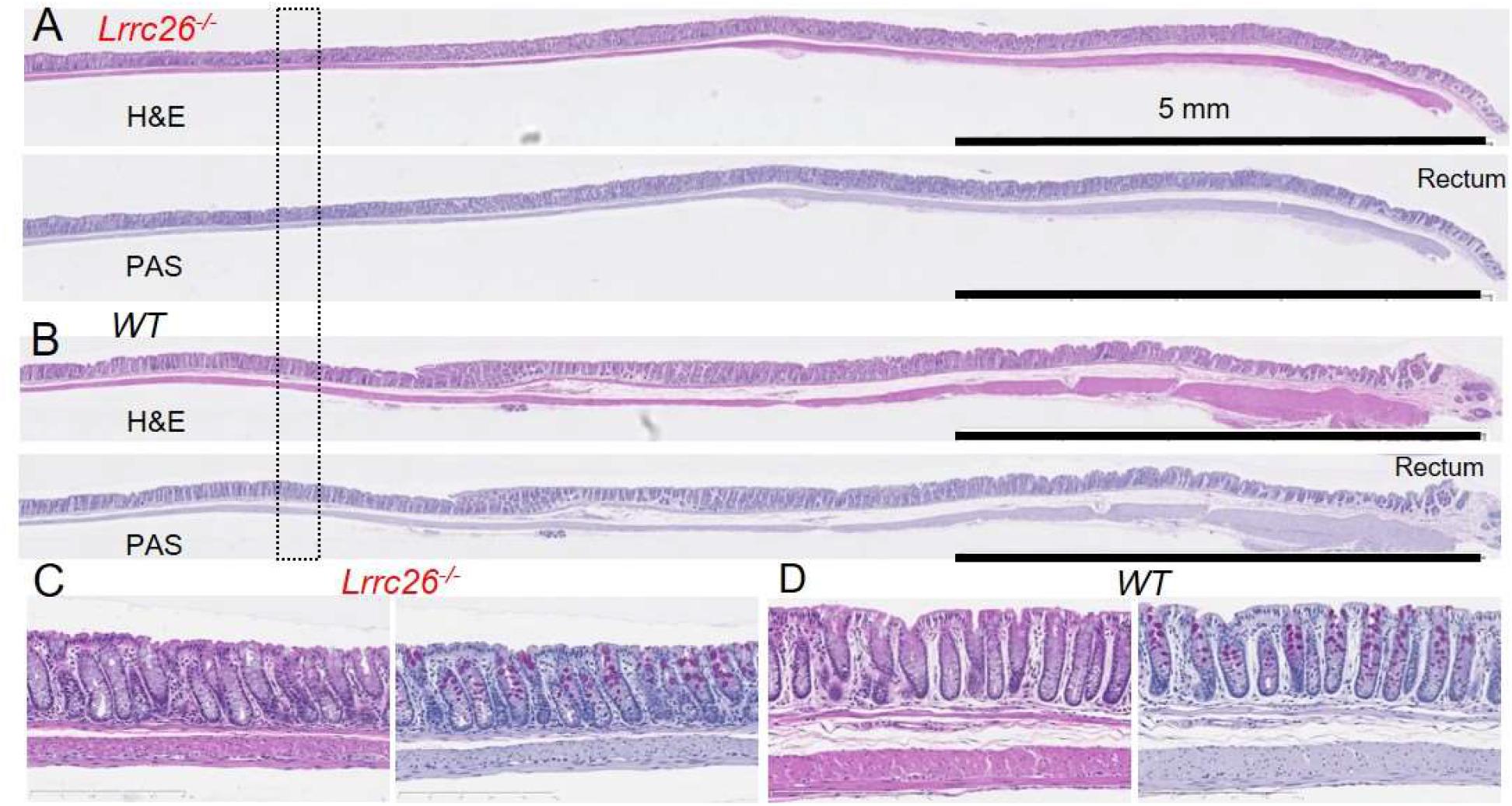
Morphological comparison between distal colons from *Lrrc26^-/-^* and *WT* littermates. **A,** Representative images of 5 micron paraffin-embedded longitudinal sections showing the entire distal third of colon from *Lrrc26^-/-^* (**A**) or *WT* (**B**) littermates, stained with hematoxylin-eosin (top) or PAS-hematoxylin (bottom). The tissues had been previously fixed with 4% formalin overnight. **C, D** Higher magnification of equivalent areas at the same level (within dotted rectangle) showing no major histological differences between genotypes.

**Figure S4.**
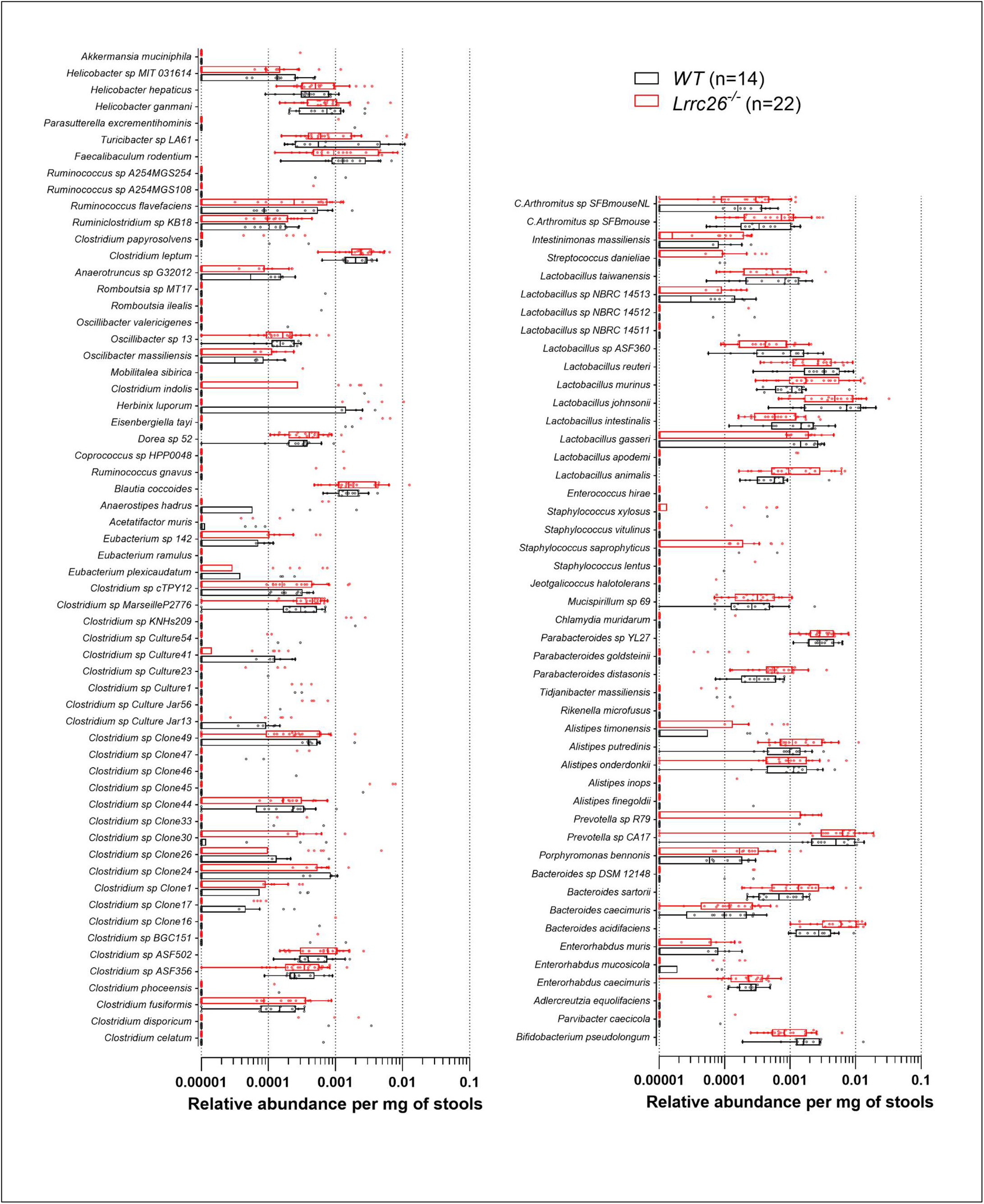
Comparison of fecal microbiota between *Lrrc26^-/-^* and *WT* mice quantified by sequencing-based analysis (16S rRNA seq). The points represents the average number of reads aligned across the different amplicons for each species in each sample, normalized by the weight of the stool sample processed. Box plot representation of the comparison in microbiota at species level between *Lrrc26^-/-^* and *WT* mice. Mice evaluated were sets of siblings belonging to 8 different litters, both sexes included, 8-12 weeks old at the time of stool collection. Stool samples were frozen immediately after collection. DNA was isolated from the stools using QIAamp Fast DNA (QIAGEN) stool mini kit. The 16S rRNA seq analysis using MRVSION computational protocol was performed by the Genome Technology Access Center at Washington University School of Medicine.

**Figure S5.**
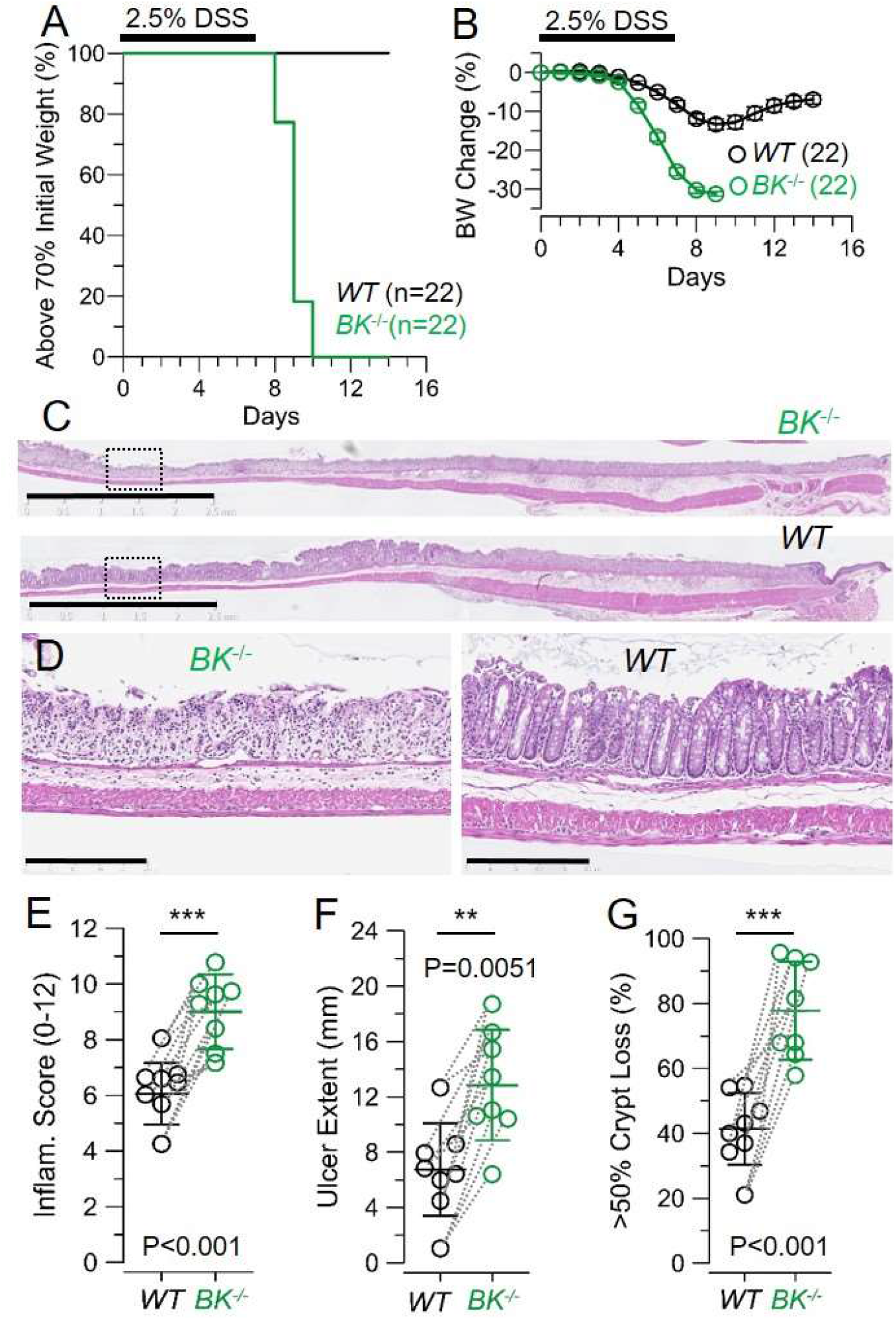
Genetic ablation of BKα subunit dramatically enhances susceptibility to DSS-induced colitis. Survival plots (**A**) and change in body weight (**B**) curves of mice treated for 7 days with 2.5% DSS (MP Biomedicals, MW: 36-50 KDa). Animals whose body weight drop > 30% of their initial weight were considered terminal and euthanized. Graphs compile results from 4 independent assays where 6 litters (with both *BK^-/-^* and *WT* littermates) Mice were 10-14 weeks old at the time of the treatment and both sexes were included. Change in body weight plot represents means±SEM. Curves were compared using a two-way ANOVA that yielded P<0.0001. **C**, Representative images of H&E stained colon tissues from *BK^-/-^* and *WT*(littermates) at the 7^th^ day of treatment, bars: 2.5 mm. **D**, Areas within the squares in (C); bars 250 μm. **E-G**, Histopathologic colitis severity in the distal colon of DSS-treated mice. Inflammation score (based on ulcer extent + crypt depletion + degree of edema)(**E**), total ulcer extention (**F**) and fraction of epithelial surface with 50% or more crypt loss (**G**). All represent means ± SD; ***, P<0.001 by two-tailed t-test.

**Fig.S6.**
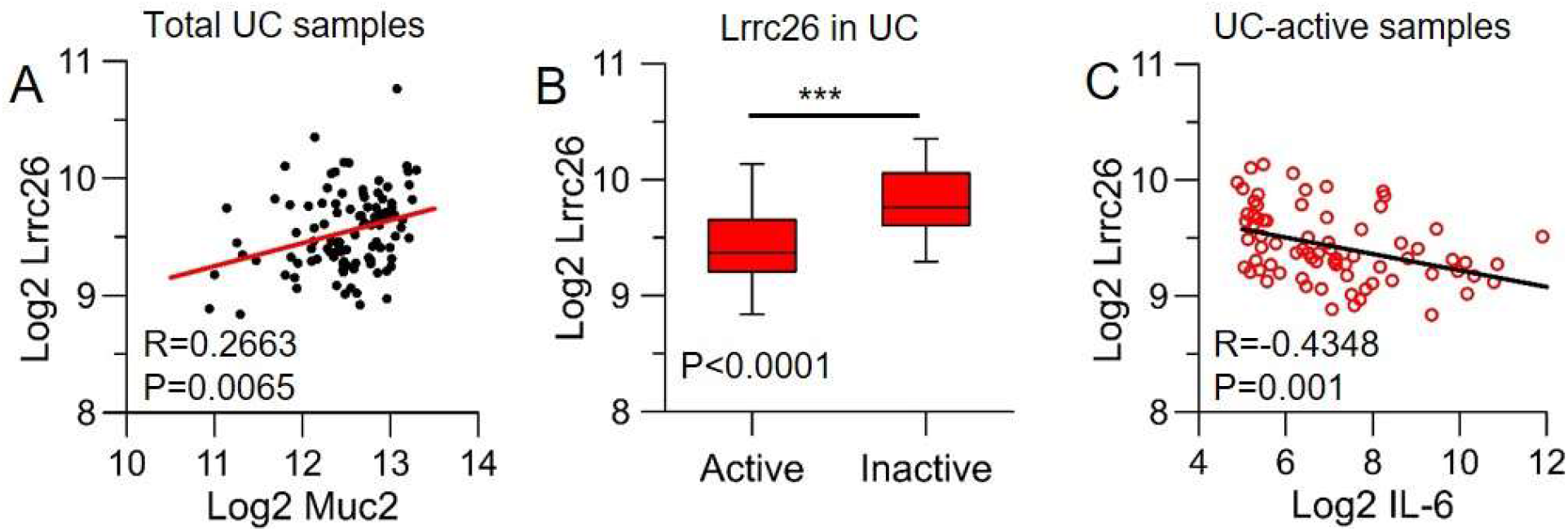
*LRRC26* gene expression profile in human samples. Public dataset GSE75214 (62), containing colon RNA expression data from healthy (Control), ulcerative colitis (UC) patients with active inflammation (Active), and UC patients without active inflammation (Inactive), was queried for expression of *LRRC26, MUC2* (as a GC marker), and *IL6* (as a marker of active inflammation). **A**, Correlation of Log2 expression of *LRRC26* and *MUC2* from all samples. **B,** The log2 expression values for *LRRC26* were grouped according to disease state. **C,** Correlation of Log2 expression of *LRRC26* and *IL6* from Active UC patients. Data represents mean ± SD. R, Pearson’s correlation coefficient; ***, P<0.001 by Mann-Whitney test.

